# Isolated human Fos promoter in plasmid DNA is overactive

**DOI:** 10.1101/2024.10.11.617784

**Authors:** Gorbunov Andrei, Fedorov Dmitrii, Kvitko Olga, Lopina Olga, Klimanova Elizaveta

**Affiliations:** Lomonosov Moscow State University, Faculty of Biology, Department of Biochemistry, Moscow, Leninskie Gory 1-12, Russia, 119234

**Keywords:** lithium, sodium, ouabain, *Fos*, transcription

## Abstract

Changes in intracellular concentrations of Na^+^ and K^+^ are shown to alter *Fos* gene expression. Here, we obtained a genetic construct encoding *TurboGFP-dest1* gene under control of the human *Fos* promoter (−549; +155) and studied its expression in HEK293T. Amplification of the *Fos* promoter sequence from genomic DNA was only efficient in the presence of Li^+^ ions. Ouabain and medium with replacement of Na^+^ with Li^+^ ions resulted in the accumulation of Na^+^ and Li^+^ in cells, respectively. These stimuli increased the mRNA level of endogenous *Fos* and the average fluorescence intensity of TurboGFP-dest1 in transfected cells. The mRNA level of *TurboGFP-dest1* was extremely higher than the mRNA level of endogenous *Fos* and was little affected by the stimuli.

## 1. Introduction

Animal cells strictly maintain gradient of Na^+^ and K^+^ across the plasma membrane mainly due to Na,K-ATPase activity. This gradient is crucial for the regulation of various cellular functions [1]. Under some physiological and pathological conditions, changes in the intracellular concentration of these monovalent cations are observed, which affects genes expression [2]. One of such genes is an early response gene *Fos*, which encodes a transcriptional factor c-Fos [3]. The basal expression of *Fos* is extremely low, however in response to external stimuli the transcription of the gene quickly increases. *Fos* gene has a complex regulation due to many regulatory sites in its composition, including serum response element (SRE), Ca^2+^/cAMP-response element (CRE), and activator protein 1 site (AP1) [4].

It was shown that an influence of the Na^+^_i_/K^+^_i_-ratio alteration on *Fos* transcription in a range of mammalian cell lines is not mediated by membrane potential, Ca^2+^-dependent signaling, intracellular pH, and cell volume changes [4–6]. This suggests the existence of an intracellular molecular sensor of monovalent cations. Several studies attempted to shed light on the nature of the Na^+^_i_/K^+^_i_-sensitive element. The first one demonstrated that SRE, CRE and AP1 elements do not activate transcription of controlled reporter gene in response to ouabain (a specific inhibitor of Na,K-ATPase) [5]. In another study, Nakagawa used molecular constructs encoding human *Fos* gene with different promoter extent to determine two sequences that are responsible for transcription activation by ouabain: the SRE-element and a region from -222 to -123 b.p. from transcription start site [7]. Remarkably, this region contains a predicted G-quadruplex sequence. G-quadruplexes are non-canonical secondary structures formed in guanine-rich regions of nucleic acids [8]. These structures bind monovalent cations, and the ability of different metal ions to stabilize G-quadruplexes typically decreases in the following order: K^+^>Rb^+^>Na^+^>Cs^+^>Li^+^ [9]. In addition, the presence of G-quadruplexes is predicted in the coding sequence and promoter of *Fos* and other Na^+^/K^+^-sensitive genes [10].

To create a tool that will further help us to study the role of G-quadruplexes in *Fos* transcription regulation, we obtained a genetic construct coding a destabilized version of TurboGFP under the *Fos* promoter control and examined the effects of ouabain and Li^+^ ions on the TurboGFP-dest1 expression in transfected HEK 293T cells. In this study, we show that lipofection causes a sodium accumulation in cells and that isolated *Fos* promoter region from -549 to +155 is overactive even without extracellular stimulation.

## 2. Materials and methods

### 2.1. Promoter-reporter constructs preparation

Reporter DNA constructs encoding the destabilized version of turboGFP protein under the control of *Fos* or *Rplp0* promoters were made. First, the *Fos* promoter region from -549 to +155 bp and the *Rplp0* promoter region from -700 to +295 bp were amplified by PCR from human HeLa cell genomic DNA using Encyclo polymerase kit from Evrogen (Russia) and primers bearing HindIII and AsiGI restriction sites (Table 1). Amplification of the *Fos* promoter was performed in a modified buffer of Chashchina [8] (60 mM Tris-HCl pH 9.5, 20 mM LiCl, 2.5 mM MgCl2, 5% DMSO). PCR products were verified by sequencing. *Fos* or *Rplp0* promoter fragments then were cloned into the promoterless peTurboGFP-PRL-dest1 vector from Evrogen (Russia) with HindIII and AsiGI restriction enzymes (SibEnzyme, Russia).

**Table 1.**
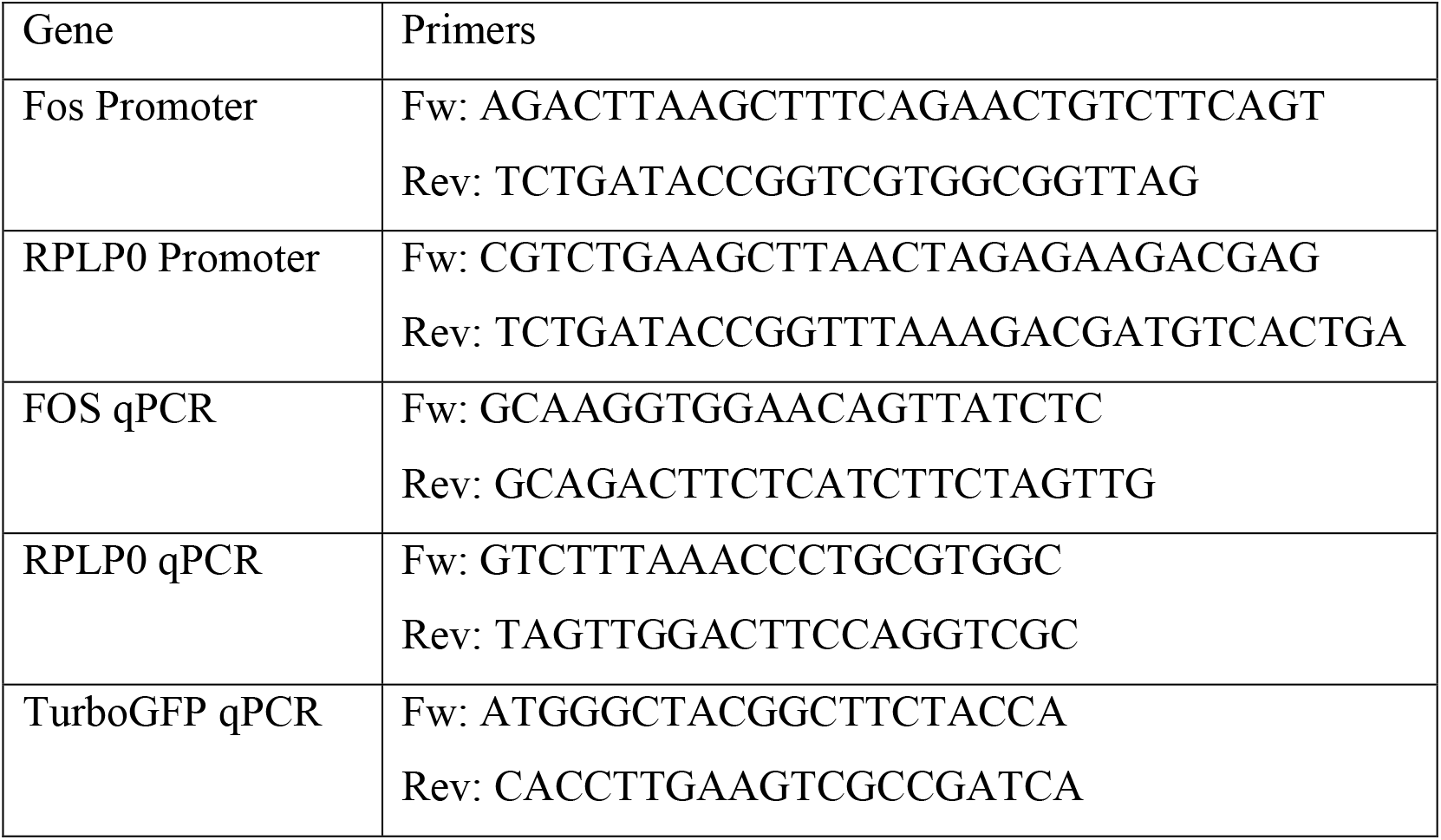
Primer sequences for promoters amplification and qPCR.

### 2.2. Cell culture and transfection protocol

HEK 293T cells were cultured in DMEM (PanEco, Russia) supplemented with 4.5 g/L glucose, 0.58 g/L glutamine (PanEco, Russia), 100 units/mL of streptomycin and penicillin (Life Technologies, USA) and 10% FBS (Cytiva, USA) at 37°C in humidified atmosphere with 5% CO_2_/balance air. Viability of cells were estimated by alamarBlue Assay (Thermo Fisher Scientific, USA) according to manufacturer instructions. Transient transfections of cells with pFOS-turboGFP or pRPLP0-turboGFP constructs were performed using Lipofectamine 2000 reagent (Invitrogen, USA). Briefly, the day before transfection, HEK 293T cells were seeded (10^5^ cells in 0.5 mL of culture medium per well) in 24-well plates pre-coated with poly-D-lysine (MP Biomedicals, USA) and cultured for 24 h. On the following day, 2 h before transfection, the culture medium was fully substituted with antibiotic-free culture medium (0.5 mL per well). HEK 293T cells were transfected with a mixture of 0.8 μg DNA and 2 μL Lipofectamine 2000 in 50 μL Opti-MEM (Gibco, USA) and incubated with DNA–liposome complexes for 24 h. The medium was then changed and cells were incubated in low-serum medium (DMEM, 4.5 g/L glucose, 0.58 g/L glutamine, 0.1% FBS) for 24 h to establish quiescence.

### 2.3. Experimental treatments

Intracellular Na^+^_i_/K^+^_i_ ratio in HEK 293T cells was altered by 3 h treatment with 1 μM ouabain (a specific inhibitor of Na,K-ATPase). In order to load HEK 293T cells with Li^+^ ions, we used a 5 h incubation with Li-medium (10 mM HEPES-LiOH pH 7.4, 130 mM LiCl, 5.2 mM KCl, 1 mM CaCl_2_, 0.5 mM MgCl_2_, 0.4 mM MgSO_4_, 0.3 mM KH_2_PO_4_, 6 mM glucose) where Li^+^ ions were used instead of Na^+^ ions. Na-medium (10 mM HEPES-NaOH pH 7.4, 130 mM NaCl, 5.2 mM KCl, 1 mM CaCl_2_, 0.5 mM MgCl_2_, 0.4 mM MgSO_4_, 0.3 mM KH_2_PO_4_, 6 mM glucose) mimicking DMEM inorganic salt composition was used as an internal control to Li-medium.

### 2.4. Determination of intracellular Na^+^, K^+^, and Li^+^ content by atomic absorption spectrometry

The ion contents in cells were determined as previously described [11] with modifications. After incubation, the plates were put on ice, the medium was discarded, and the cells were washed three times with 3 mL of ice cold 0.1 M MgCl_2_ solution. Then, 1.5 mL of 5% trichloroacetic acid (TCA) was added to each well and incubated for several hours at 4°C, after which the contents were scraped off and transferred to Eppendorf tubes. The obtained samples were centrifuged for 10 min at 18,000 g. The supernatant was collected in individual tubes, while the residual TCA was aspirated, after which the precipitate was dissolved in 0.1 M NaOH, 0.1% sodium deoxycholate solution and protein concentration in it was determined by Lowry method [12]. Contents of Na^+^, Li^+^ and K^+^ in the TCA extracts were measured by flame atomic absorption spectrometry with a Kvant-2m1 spectrometer (Kortek, Russia) in a propane-air mixture at wavelengths of 589,6 nm, 670,8 nm И 766,5 nm, respectively. NaCl (0.05–2 mg/liter Na^+^), LiCl (0.05–2 mg/liter Li^+^) and KCl (0.5–2 mg/liter K^+^) solutions containing 5% TCA were used as references. Contents of Na^+^_i_, K^+^_i_, and Li^+^_i_ in each well were normalized to the amount of protein in the same well.

### 2.5. Live-cell fluorescent imaging and image analysis

HEK 293T cells were cultured and transfected with pFOS-turboGFP or pRPLP0-turboGFP as described in p. 2.2 and then subjected to the treatment specified in p. 2.3 a certain number of hours before the end of incubation for 24 hours in a culture medium containing 0.1% FBS. Cells were imaged inside Tokai hit (Japan) gas chamber at 37 °C in humidified atmosphere with 5% CO_2_/balance air using an inverted Olympus confocal microscope (Japan). Fluorescent signals of turboGFP-dest1 were acquired from a single optical plane (pinhole diameter = 130 μm) via Olympus 10x (NA 0.4) objective using Olympus software. TurboGFP-dest1 was excited by 488 nm laser line, and its fluorescence was acquired within the spectral range 490–540 nm. The resulting images were processed using Fiji software. Individual cells were selected, when critical intensity level, visually separating cells from the background in the best way, was set. The average fluorescence intensity was calculated inside each selected cell.

### 2.6. Analysis of gene transcription by real-time polymerase chain reaction (RT-PCR)

Plates were transferred on ice, HEK 293T cells were washed with 1 mL of ice-cold Hanks’ solution without Ca^2+^, Mg^2+^ salts and phenol red, and 400 μl of Trizol reagent (Invitrogen, USA) per cell were added for total RNA isolation. After separation of the aqueous phase containing nucleic acids using chloroform and treatment with 96% ethanol, further steps of RNA isolation and DNAase treatment were performed on Qiagen RNeasy kit columns (Neitherlands). An ImProm-IITM Reverse Transcription System kit (Promega, USA) was used for the reverse transcription reaction according to the manufacturer’s instructions based on 0.5 μg of total RNA from each probe. Real-time PCR was performed using the Bio-Rad Real-Time PCR System (BioRad, USA). Amplification protocol: 1) 95°C 5 min; 2) 95°C 10 s; 3) 58°C 10 s; 4) 72°C 20 s (40 repetitions of steps 2-4). Primers (Table 1) were designed using the NCBI and BLAST searchable databases and synthesized by Syntol (Russia). Expression level of each gene of interest was calculated using the *Rplp0* as a reference gene by ΔΔCt method [13]. Average gene expression of *Fos* gene in the control samples was taken as 100%. PCR products were verified by sequencing.

### 2.7. Statistical data analysis

Statistical data analysis was performed using the Origin software package (OriginLab Corporation, USA). All sample distributions were checked for normality using the Shapiro-Wilk test. To compare two independent groups, the Student’s t-test was used for normally distributed data, as well as the Mann-Whitney U-test for non-normally distributed samples. Multiple comparisons for normally distributed data were performed using single-factor analysis of variance (ANOVA) followed by the Tukey test.

## 3. Results and discussion

To investigate the mechanism of Na^+^_i_/K^+^_i_-dependent gene transcription regulation we obtained molecular constructs encoding TurboGFP-dest1 (Fig. 1A) under human *Fos* promoter control (from -549 to +155 b.p. from transcription start site, pFOS-TurboGFP) or human *Rplp0* promoter control (from -700 to +295 b.p., the sequence does not have any predicted G4 structures, pRPLP0-TurboGFP). To do this we amplified *Fos* and *Rplp0* promoter sequences from genomic DNA of HeLa cells. First of all, *Rplp0* promoter was successfully amplified using a commercial Encyclo PCR buffer, whereas *Fos* promoter was not. In a buffer for GC-rich sequences *Fos* promoter was amplified but with a deletion in 78 b.p (Fig. 1B). We suppose that it is linked to G4-structures in *Fos* sequence. G4 structures are known to interfere PCR in the presence of K^+^ ions. However, most of commercial buffers for DNA-polymerases contain K^+^ ions. To overcome this, we used a modified buffer of Chashchina [8] with Cs^+^ replaced by Li^+^ ions. Using it, we successfully amplified full-length sequence of *Fos* promoter.

**Figure 1.**
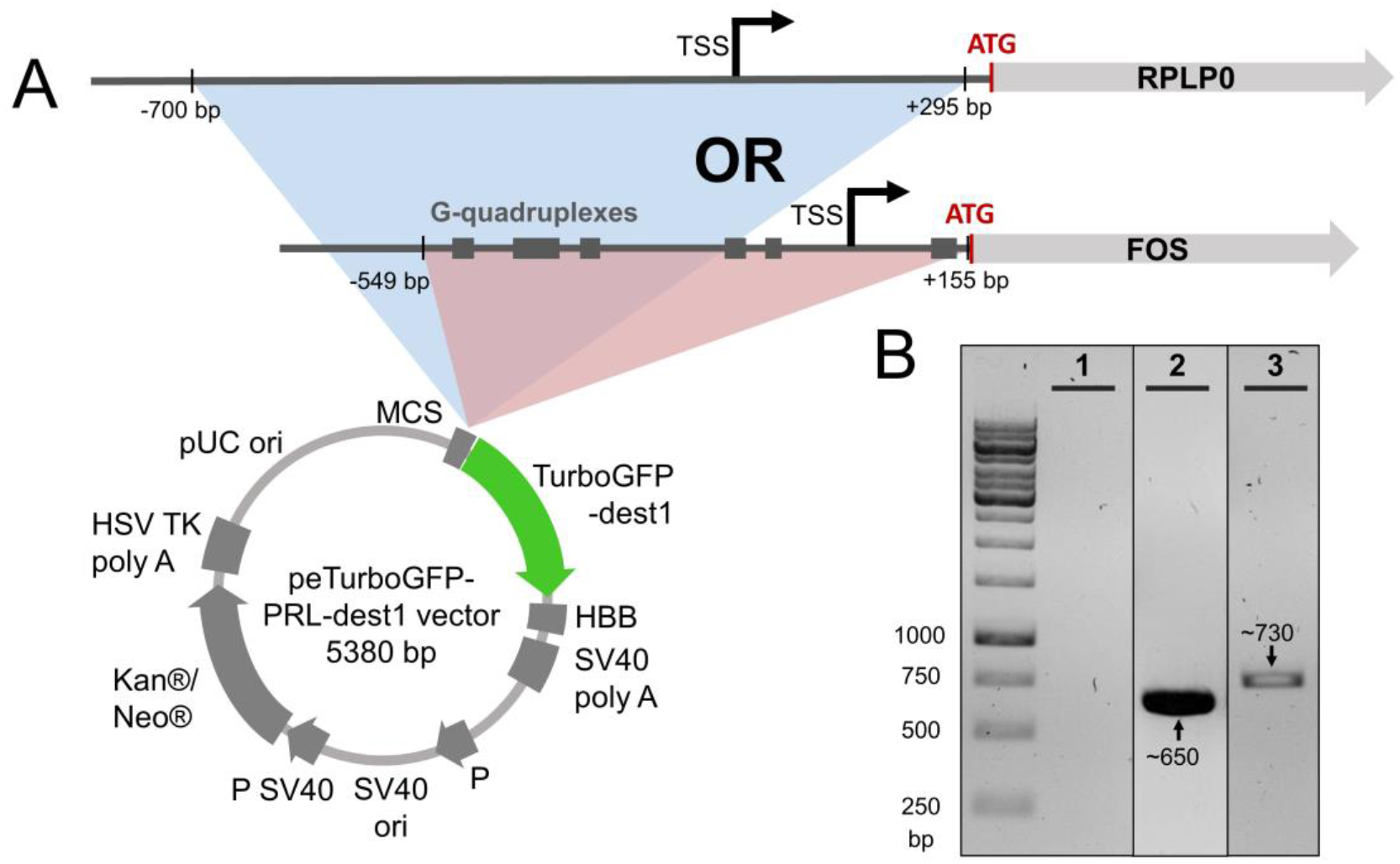
Reporter DNA vectors used to study the activity of the FOS gene promoter region. **(A)** Schematic representation of the pFOS-turboGFP and pRPLP0-turboGFP vectors encoding the destabilized version of turboGFP protein under the control of the promoter region of *Fos* (from -549 to +155 bp) or *Rplp0* (from -700 to +295 bp) genes. **(B)** Agarose gel (1,5%) electrophoresis of PCR products of the *Fos* gene promoter region from -549 to +155 bp obtained in different PCR buffers: commercial buffers for GC-rich sequences (1) and Encyclo buffer (2) or a special buffer where K^+^ ions were substituted with Li^+^ ions (3). The amplified DNA was designed for subsequent cloning and estimated length of the PCR product (including added restriction sites) was 728 bp.

The next step was to evaluate an influence of transfection procedure on ion composition of HEK 293T cells. Fig. 2A shows that lipofection with plasmid DNA pFOS-TurboGFP or pRPLP0-TurboGFP increases intracellular Na^+^ content by 40-60% 24 h after transfection, whereas lipofection without DNA does not affect this parameter. At least two factors may cause such an impact. First, ion transport may be affected through the RNA polymerase III dependent pathway of interferon expression induction by foreign AT-rich dsDNA [14]. Another factor is an expression of TurboGFP-dest1 protein and its effects on cellular functions. TurboGFP-dest1 is known to interfere with NF-kB pathway in T cells, HEK293, and HeLa cells [15], thus it is possible that the protein may alter ion transport. However, the exact mechanisms remain to be determined.

**Figure 2.**
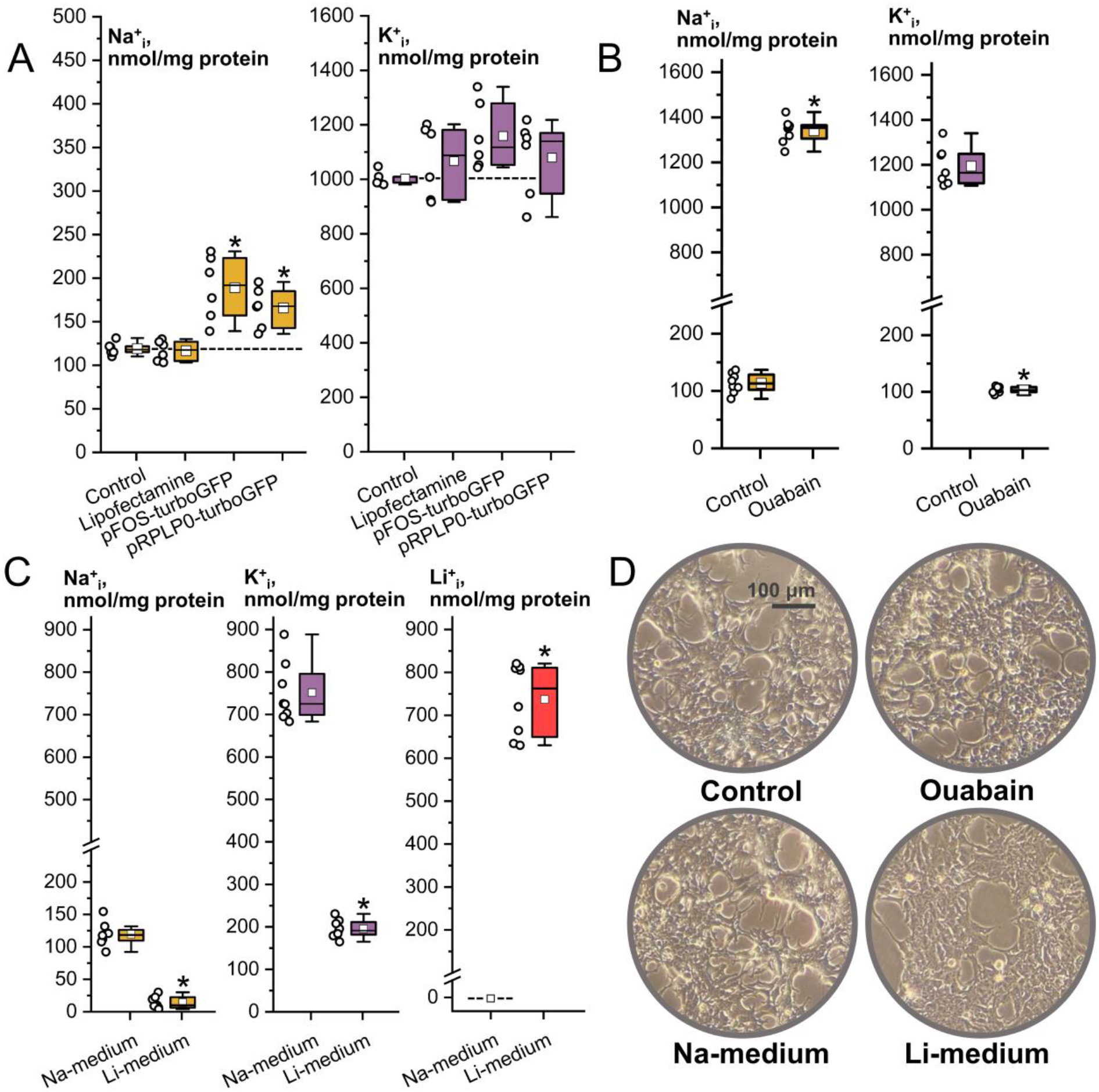
Effect of different experimental treatments on the intracellular content of monovalent metal cations in HEK 293T cells. **(A)** Transfection of HEK 293T cells with FOS-turboGFPp and RPLP0-turboGFPp increases Na^+^_i_ content by 60% and 40%, respectively, and has no significant effect on K^+^_i_. Transfection reagent Lipofectamine 2000 itself does not affect neither Na^+^_i_ nor K^+^_i_. **(B)** Treatment of these cells with 1 μM ouabain for 3 h dramatically increases intracellular Na^+^ content almost 12-fold and has the opposite effect on K^+^_i_. **(C)** HEK 293T cells incubated in Li-medium (with 135 mM of Li^+^ and 5,5 mM of K^+^) for 5 h accumulate Li^+^ ions up to 740 nmol/mg of protein and show significantly lower Na^+^_i_ and K^+^_i_ contents compared to Na-medium (135 mM Na^+^ and 5,5 mM K^+^) – control-type medium which represents DMEM in Na^+^ and K^+^ content. **(D)** Photographic images of HEK 293T cells exposed to 1 μM of ouabain for 3 h or incubated in Na-or Li-media for 5 h. Data are shown as experimental points (n=6-8) along with boxplot diagrams (with whiskers spreading for 1,5 interquartile ranges). Statistical analysis was performed using t-test or ANOVA followed by post hoc Tukey test (* -p < 0,05).

To compare the effects of Li^+^ and Na^+^ ions on *Fos* promotor activity in transfected HEK 293T, cells were incubated in DMEM with 0.1% FBS in the presence of 1 µM ouabain or in a medium containing 135 mM Li^+^ (Li-medium). Fig. 2B shows that incubation of HEK 293T cells with 1 µM ouabain for 3 h led to a decrease in K^+^_i_ content of 11.5 times and an increase in Na^+^_i_ content of 11.8 times. An incubation with Li-medium for 5 h resulted in 740 nmol Li^+^/mg of protein being accumulated in HEK 293T cells as well as in a decrease of Na^+^_i_ and K^+^_i_ content by 85% and 90%, respectively (Fig. 2C). Both stimuli did not change cell viability (data not shown), but Li-medium slightly affected cell morphology (Fig. 2D).

To estimate *Fos* promoter activity in transfected cells, we evaluated a fluorescence intensity of TurboGFP-dest1 by confocal microscopy and TurboGFP-dest1 transcription level by RT-PCR. Fig. 3 shows that medians of fluorescence intensity in cells transfected with pFOS-TurboGFP were increased by 21% and 11% after incubation with 1 µM ouabain and in Li-medium, respectively, whereas in case of cells transfected with pRPLP0-TurboGFP we did not observe significant changes in fluorescence intensity. The fluorescence intensity values were very close for cells transfected with pFOS-TurboGFP and pRPLP0-TurboGFP plasmids. RT-PCR data showed no significant changes in TurboGFP-dest1 mRNA level, whereas endogenous *Fos* mRNA level was increased by 17 and 10 times in cells incubated with ouabain and in Li-medium, respectively (Fig. 4). TurboGFP-dest1 mRNA level was 1000-3000 times larger than endogenous *Fos* mRNA level in all experimental and control samples. We suggest that these results indicate an excessive activation of *Fos* promoter in pFOS-TurboGFP construct, whose activity is close to housekeeping genes such as *Rplp0*. Probably due to this, an effect of experimental treatments on *Fos* promoter activity in the vector is negligible.

**Figure 3.**
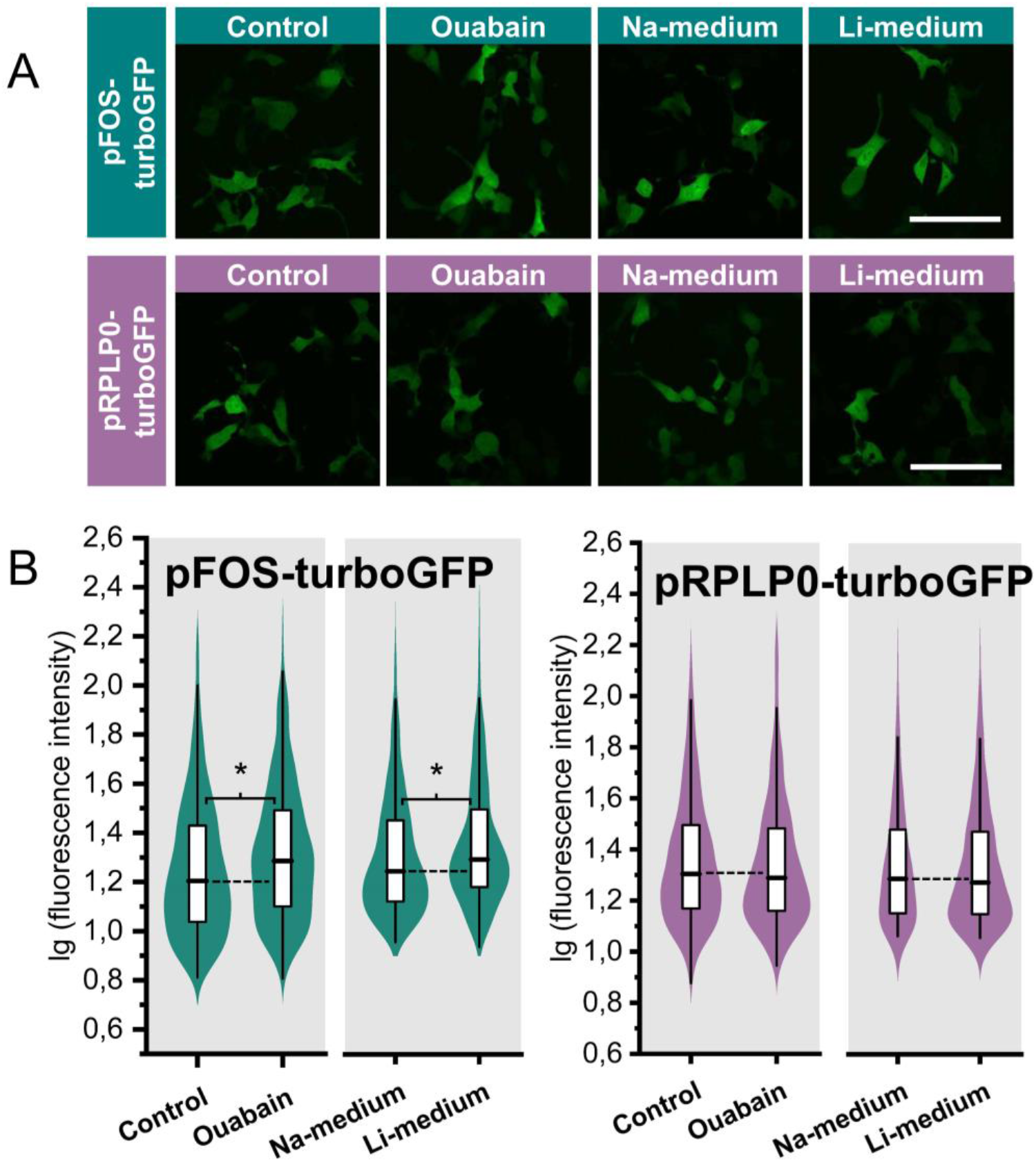
Expression of turboGFP-dest1 reporter protein under the control of *Fos* or *Rplp0* promoters in HEK 293T cells. **(A)** Confocal fluorescent images of HEK 293T cells transfected with pFOS-turboGFP or pRPLP0-turboGFP constructs. Two days after transfection, cells were treated with 1 μM ouabain for 3 h or were incubated in Na-or Li-media for 5 h. Scale bars: 100 μm. **(B)** Distributions of mean intensity of turboGFP-dest1 fluorescence in single cells (n=500 – 1000). Statistical analysis revealed low but significant elevation of the reporter protein fluorescence in response to treatment of HEK 293T cells with ouabain vs. control (DMEM). The same effect was shown in cells incubated in Li-medium compared to Na-medium. Statistical analysis was performed using Mann–Whitney U test (* - p < 0,05).

**Figure 4.**
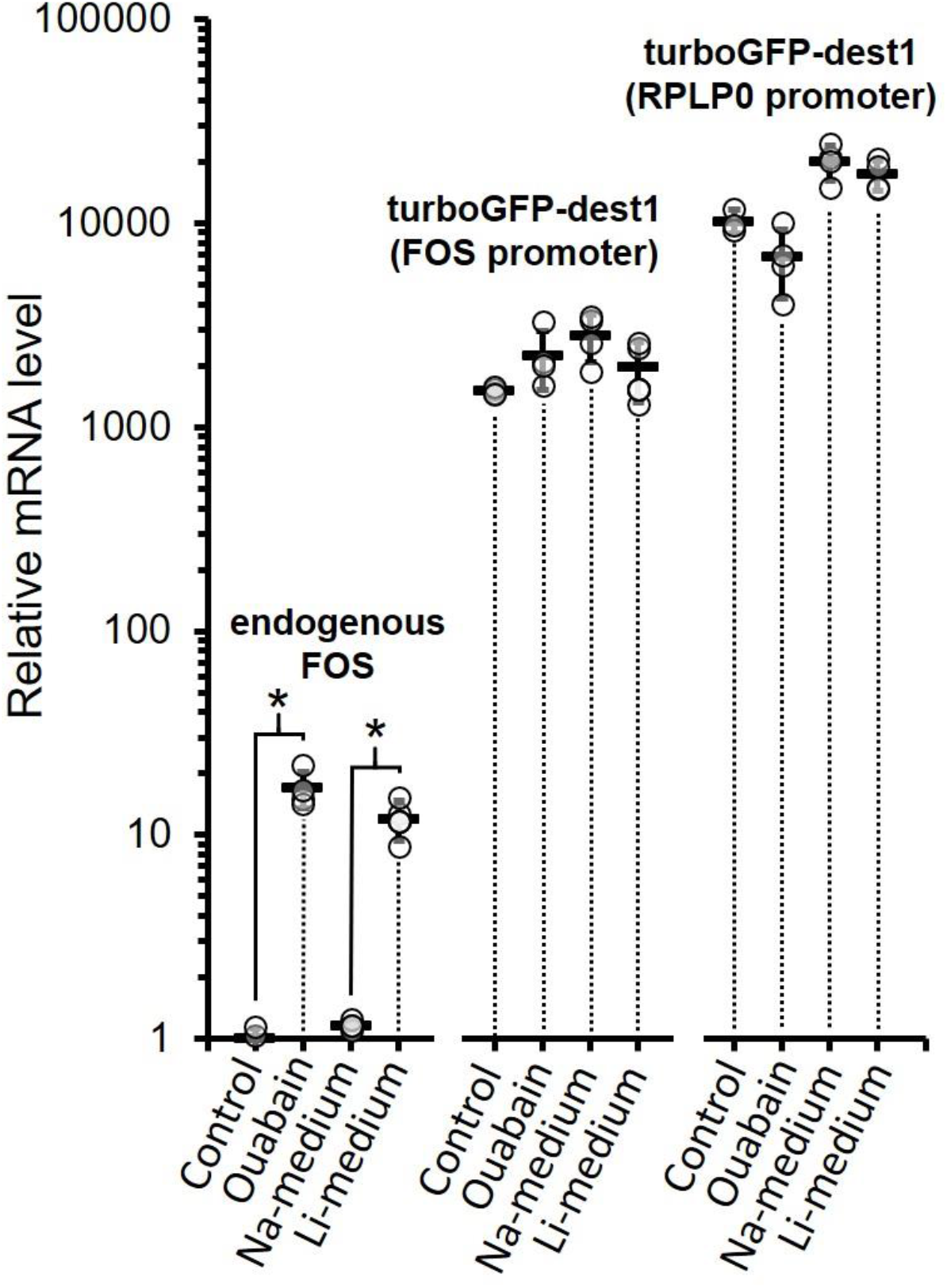
Relative expression levels of endogenous *Fos* mRNA or turboGFP-dest1 mRNA under the control of *Fos* or *Rplp0* promoter in HEK 293T cells. HEK 293T cells were transfected with pFOS-turboGFP or pRPLP0-turboGFP. Two days after transfection, cells were treated with 1 μM ouabain for 3 h or were incubated in Na-or Li-media for 5 h. Cell lysates then were analyzed for endogenous *Fos* or turboGFP-dest1 mRNA levels using RT PCR technique. Data are shown as mean ± standard deviation (SD) along with experimental points (n=4). Statistical analysis revealed prominent and significant elevation of endogenous FOS mRNA in response to ouabain or incubation in Li-medium vs. control or Na-medium incubation, respectively. Statistical analysis was performed using t-test, * -p < 0,05).

An effect of Na^+^_i_/K^+^_i_ ratio change on *Fos* promoter activity was studied before. A construct containing *firefly* luciferase gene under control of *Fos* 5’-UTS from -1264 to +103 b.p. was made. The luciferase expression from the construct was not affected by ouabain [4], which is similar to our results. In another study, a construct based on full-length human *Fos* gene was used. The results of the study show significant activation of *Fos* transcription in the construct upon ouabain treatment. Summing these results with ours, we suggest that regulatory elements in coding region of *Fos* gene may deactivate transcription at rest. It has been shown that a strong transcription elongation block is located within the first intron of the *Fos* gene (in the region from +363 to +387 bp) [16–19]. This regulatory region represses basal expression of not only the *Fos* promoter, but also the expression of the “strong” metallothionein-2 gene promoter in a chimeric construct where the *Fos* open reading frame was placed under the control of this promoter [18]. In addition, there are other possible explanations of our results: an absence of far regulatory elements in obtained vector or differences between modification patterns of plasmid DNA and genomic DNA. However, these assumptions require further confirmation.

## Conclusion

We demonstrated *Fos* upregulation in HEK 293T cells by ouabain and intracellular Li^+^ accumulation. We also generated a genetic construct with TurboGFP-dest1 under the control of the *Fos* promoter. TurboGFP-dest1 expression in HEK293T cells transfected with the construct was abnormally high and almost insensitive to changes in the intracellular ionic composition. It is likely that other regulatory elements outside the studied promoter region are required to inhibit *Fos* expression in the resting state. Another conclusion is that lipofection of HEK 293T cells with genetic constructs expressing TurboGFP-dest1 leads to Na^+^ accumulation in the cells. This phenomenon may be due to the activation of the intracellular response system to foreign DNA or a nonspecific effect of TurboGFP-dest1 on ion transport.

## Abbreviations

AP1,: activator protein 1;
CRE,: Ca^2+^/cAMP-response element;
DMEM,: Dulbecco’s modified Eagle medium;
DMSO,: dimethyl sulfoxide;
FBS,: fetal bovine serum;
HEPES,: 4-(2-hydroxyethyl)-1-piperazineethanesulfonic acid;
NA,: numerical aperture;
pFOS-turboGFP,: a genetic construct encoding *TurboGFP-dest1* gene under control of the human *Fos* promoter;
pRPLP0-turboGFP,: a genetic construct encoding *TurboGFP-dest1* gene under control of the human *Rplp0* promoter;
RT-PCR,: reverse transcription-polymerase chain reaction;
SRE,: serum response element;
TCA,: trichloroacetic acid;
5’-UTS,: 5’-untranslated sequence.

## Ethics declarations

This article does not contain any studies with human participants or animals performed by any of the authors.

## Data availability statement

Data will be made available on request.

## CRediT authorship contribution statement

Gorbunov Andrei: Investigation, Writing – original draft, Formal Analysis, Visualization. Fedorov Dmitrii: Investigation, Writing – original draft. Kvitko Olga: Investigation. Lopina Olga: Writing – review & editing, Conceptualization. Klimanova Elizaveta: Writing – review & editing, Conceptualization, Funding acquisition.

## Acknowledgments

This study was supported by Russian Science Foundation, grant number 19-75-10009.

